# Hierarchical modeling of haplotype effects based on a phylogeny

**DOI:** 10.1101/2020.01.31.928390

**Authors:** Maria Lie Selle, Ingelin Steinsland, Finn Lindgren, Vladimir Brajkovic, Vlatka Cubric-Curik, Gregor Gorjanc

## Abstract

This paper introduces a hierarchical model to estimate haplotype effects based on phylogenetic relationships between haplotypes and their association with observed phenotypes. In a population there are usually many, but not all possible, distinct haplotypes and few observations per haplotype. Further, haplotype frequencies tend to vary substantially - few haplotypes have high frequency and many haplotypes have low frequency. Such data structure challenge estimation of haplotype effects. However, haplotypes often differ only due to few mutations and leveraging these similarities can improve the estimation of haplotype effects. There is extensive literature on this topic. Here we build on these observations and develop an autoregressive model of order one that hierarchically models haplotype effects by leveraging phylogenetic relationships between the haplotypes described with a directed acyclic graph. The phylogenetic relationships can be either in a form of a tree or a network and we therefore refer to the model as the haplotype network model. The haplotype network model can be included as a component in a phenotype model to estimate associations between haplotypes and phenotypes. The key contribution of this work is that by leveraging the haplotype network structure we obtain a sparse model and by using hierarchical autoregression the flow of information between similar haplotypes is estimated from the data. We show with a simulation study that the hierarchical model can improve estimates of haplotype effects compared to an independent haplotype model, especially when there are few observations for a specific haplotype. We also compared it to a mutation model and observed comparable performance, though the haplotype model has the potential to capture background specific effects. We demonstrate the model with a case study of modeling the effect of mitochondrial haplotypes on milk yield in cattle.

## 1 Introduction

This paper develops a hierarchical model to estimate haplotype effects based on phylogenetic relationships between haplotypes and their association with observed phenotypes. With current technology we can readily obtain genome-wide information about an individual, either through single-nucleotide polymorphism array genotyping or sequencing platform. Since the genome-wide information has become abundant, modeling this data has become the standard in animal and plant breeding as well as human genetics. The application of this modeling has been shown to improve genetic gains in breeding (Meuwissen et al., 2001; Hickey et al., 2017; Ibanez-Escriche and Simianer, 2016) and has potential for personalized prediction in human genetics and medicine (Begum, 2019; de los Campos et al., 2018; Lello et al., 2018; Maier et al., 2018).

The aim of geneticists is to infer which mutations are causing variation in phenotypes and what are their effects. This aim is nowadays approached with genome-wide association studies of regressing observed phenotypes on mutation genotypes (see the recent review of Morris and Cardon, 2019). However, mutations arise on specific haplotypes passed between generations, which limits accurate estimation due to low frequency of mutations, correlation with other mutations and limited ability to observe all mutations with a used genomic platform (e.g., see Gibson, 2018; Simons et al., 2018; Uricchio, 2019). Further, most mutations do not affect a phenotype, while some mutations have background (haplotype) specific effects (e.g., Chandler et al., 2017; Wojcik et al., 2019; Steyn et al., 2019).

Instead of focusing on mutation effects we here focus on haplotype effects and their differences to estimate the effect of mutations on specific haplotypes. There is extensive literature on estimating haplotype effects (Balding, 2006; Thompson, 2013; Morris and Cardon, 2019). One issue with estimating haplotype effects is that there is usually an uneven distribution of haplotypes in a population (Ewens, 1972, 2004; Walsh and Lynch, 2018) and estimating the effects of rare haplotypes is equally challenging as estimating the effect of rare mutations. However, the described genetic processes in the previous paragraph create a “network” of haplotypes (sometimes referred to as *genealogy* or *phylogeny*), which suggests that effects of similar haplotypes are similar. This observation inspired Templeton et al. (1987) to cluster phylogenetically similar haplotypes. Others have used similar approaches to utilize this data structure (Balding, 2006; Thompson, 2013; Morris and Cardon, 2019).

We here approach the problem of estimating haplotype effects by leveraging phylogenetic relationships between haplotypes described with a directed acyclic graph (DAG) and developing a hierarchical model of haplotype effects on this graph. We were inspired by recent advances in building phylogenies on large datasets (Kelleher et al., 2019) and aimed to develop a hierarchical model that could scale to a large number of haplotypes. Our work extends the phylogenetic mixed modeling of the whole genome (Lynch, 1991; Pagel, 1999; Housworth et al., 2004; Hadfield and Nakagawa, 2010) to a specific region. This region specific modeling could be applied either across species (macroevolution) or within a species (microevolution).

A potentially important modeling aspect with respect to across and within species modeling is that the phylogenetic mixed model assumes Brownian motion for evolution of phenotypes along a phylogeny (Felsenstein, 1988; Huey et al., 2019). Brownian motion is a continous random-walk process with variance that grows over time (is non-stationary) (Blomberg et al., 2019; Gardiner, 2009), which makes it a plausible model of evolution due to mutation and drift. There are alternatives to Brownian motion, in particular the Ornstein-Uhlenbeck process that can accomodate various forms of selection (Lande, 1976; Hansen and Martins, 1996; Martins and Hansen, 1997; Paradis, 2014). Ornstein-Uhlenbeck process is also a continuous random-walk, but with an additional parameter that reverts the process to the mean (is a stationary process; (e.g., Gardiner, 2009; Blomberg et al., 2019)). Both of these models imply Gaussian distribution for the initial state and increments. The differences between the two processes might be important in the context of modeling haplotypes that likely manifest less variation than whole genomes, particularly when considering haplotypes within a species or even specific population.

The aim of this paper is to develop a hierarchical model for haplotype effects by leveraging phylogenetic relationships between haplotypes. We assume that such relationships are encoded with a DAG and therefore call the model the haplotype network model. Since haplotypes differ due to a small number of mutations and very few mutations have an effect we expect that phylogenetically similar haplotypes will have similar effects. Furthermore, the small discrete number of mutation differences suggest discrete-time analogues of Brownian and Ornstein-Uhlenbeck processes. Therefore, we have modeled the effect of a mutated haplotype given its parental haplotype with a stationary autoregressive model of order one following the phylogenetic structure encoded with a DAG. The results show that the haplotype network model improves the estimation of haplotype effects compared to an independent haplotype model due to sharing of information. The results also show that it is comparable to a mutation model, but as we discuss it has a potential to capture background specific effects.

## 2 Material and Methods

In this section we present the haplotype network model and show how to use it as a component in a phenotype model. We also describe simulations, a case study of modeling mitochondrial effects on milk yield in cattle, and the chosen method to perform inference and model evaluation.

### 2.1 The haplotype network model

Here, we present the haplotype network model, which is a hierarchical model for haplotype effects based on phylogenetic relationships between haplotypes encoded with a directed acylic graph. The phylogenetic relationships can be either in a form of a tree or a more general network. We also present two generalizations of the model - first due to multiple parental haplotypes and second due to genetic recombination.

We assume throughout that phylogeny between haplotypes is known and that it can be encoded with a DAG. The haplotype network model can in principle deal with different types of mutations, but for simplicity we focus only on biallelic mutations with the code 0 used for the ancestral/reference allele (commonly at a higher frequency in a population) and the code 1 used for the alternative allele that arose due to a mutation.

#### 2.1.1 Motivating example

To motivate the haplotype network model, we use the example from Kelleher et al. (2019) that presents 5 haplotypes spanning 7 bialelic polymorphic sites (Table 1). Note that the 5 haplotypes are just a sample of the 2^7^ = 128 possible haplotypes over the 7 sites. An example of a phylogeny for the haplotypes is shown in Figure 1, where haplotypes are denoted as nodes (we also show their allele sequence), relationships between haplotypes are denoted as edges and mutated sites are denoted with a number on edges. For example, the ancestral haplotype *i* has allele sequence 0000000, and the haplotype *g* with sequence 1000100 differs from the ancestral haplotype due to mutations at the sites 5 and 1.

**Table 1:**
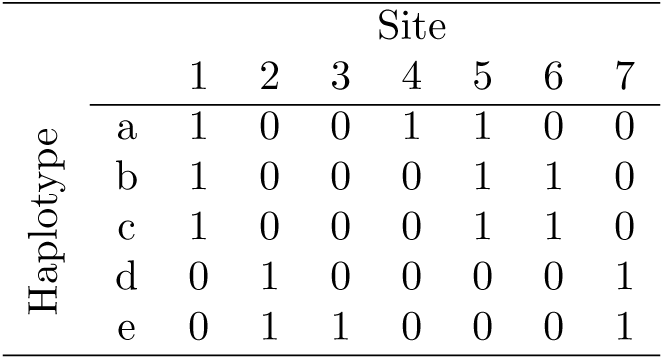
Example of 5 haplotypes spanning 7 mutations from Kelleher et al. (2019). The ancestral (reference) alleles are coded as 0 and alternative alleles are coded as 1

**Figure 1:**
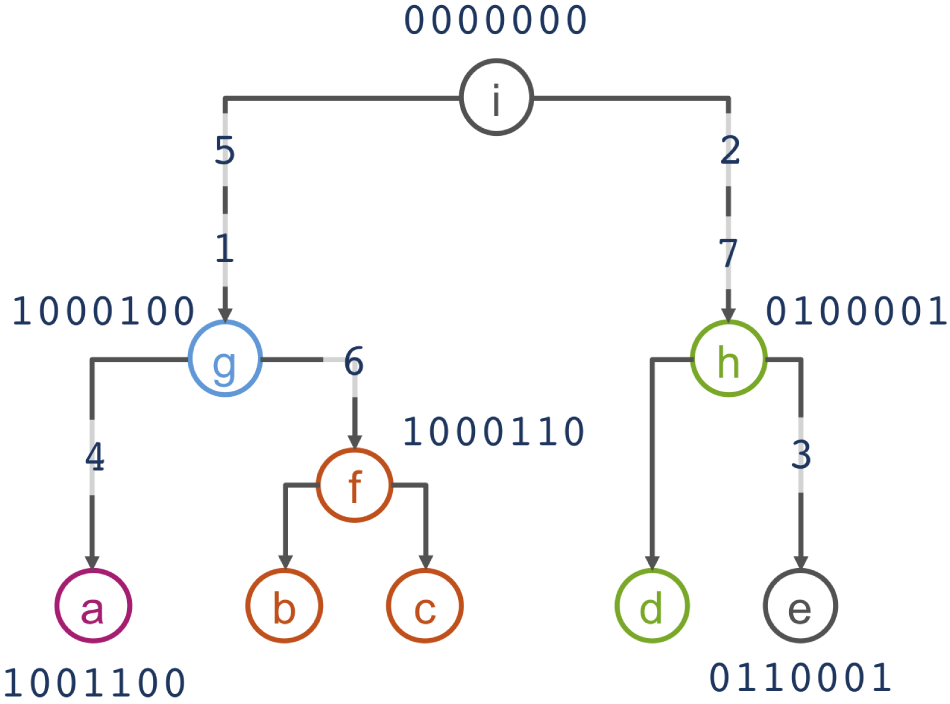
Phylogenetic relationship of haplotypes in Table 1

Assuming that similar haplotypes have similar effects we model dependency between parent-progeny pairs of haplotypes with an autoregressive Gaussian process of order one. For haplotypes in Table 1 and Figure 1 this model implies the following set of conditional dependencies:

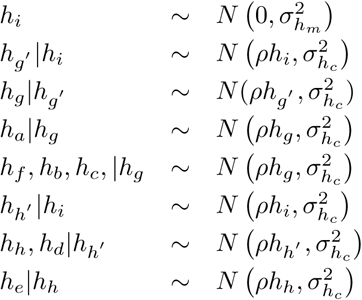

where *h*_*i*_, *h*_*g*_, …, *h*_*e*_ indicate the effect of haplotypes *i, g*, …, *e* and 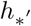 indicates the effect of haplotypes that occur between haplotypes separated by multiple mutations, for example, *g*′ is the additional haplotype between the haplotypes *i* and *g* due to two mutations between *i* and *g*; we describe the other model parameters 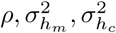 in the following.

#### 2.1.2 The model

Assume a known general phylogenetic network of haplotypes described with a DAG (DAG) with haplotype effects as nodes and relationships between the haplotype effects as edges as in Figure 1. We model the effect of a chosen “starting” (central, ancestral, most common, etc.) haplotype 0 with mean-zero and marginal variance 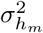:

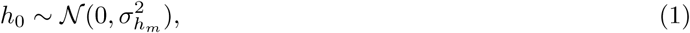

and any other haplotype *j* in the phylogenetic network as a function of its one-mutation-removed parental haplotype *p* (*j*) assuming the autoregressive Gaussian process of order one with the autocorrelation between haplotype effects of *ρ* (|*ρ*| < 1) and conditional variance of 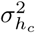:

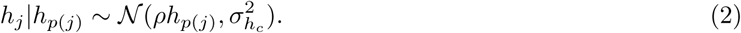

We consider the autoregressive Gaussian process of order one that is stationary both in mean and variance, which is achieved by setting the marginal variance to 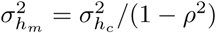, so 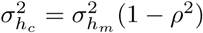. This is the standard autoregressive model of order one used in time-series analysis (Rue and Held, 2005). The difference here is that we are applying the model onto a phylogenetic network described with a DAG.

The set of distributions in (1) and (2) give a system of equations for all *n* haplotype effects ***h*** = (*h*_1_, …, *h*_*n*_)^*T*^ :

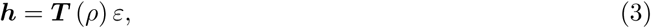

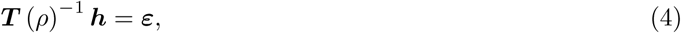

where the matrices ***T*** (*ρ*) and ***T*** (*ρ*)^−1^ respectively represent marginal and conditional phylogenetic regression between haplotype effects ***h*** and the vector ***ε*** represents haplotype effect deviations, 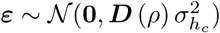. The expression ***T*** (*ρ*) indicates that the matrix ***T*** depends on the value of *ρ*. Since haplotype effect deviations are independent the matrix ***D*** (*ρ*) is diagonal and has value 1*/*(1 − *ρ*^2^) for the “starting” haplotype and 1 for the other haplotypes. Following the assumed autoregressive process of order one (2), the non-zero elements of ***T*** (*ρ*)^−1^ are 1 along the diagonal and −*ρ* between a haplotype effect (row index) and its parental haplotype effect (column index). This simple sparse lower-triangular structure of the matrix ***T*** (*ρ*)^−1^ arises from the Markov properties of the autoregressive process (Rue and Held, 2005).

From (3) covariance between haplotype effects are:

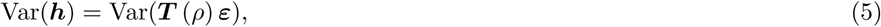

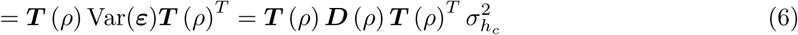

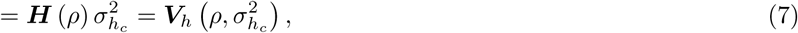

The covariance expression (5) shows that haplotype covariances 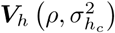 depend on the autocorrelation and variance parameters, while the covariance coefficients ***H*** (*ρ*) depend only on the autocorrelation parameter. So, the variance parameter is capturing scale (spread) of effects and the autocorrelation parameter is capturing the dependency structure. Note that these two parameters are correlated by definition 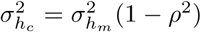. When *ρ* = 0 there is no covariance between haplotype effects due to phylogenetic relationships, which suggests a model where haplotype effects are identically and independently distributed, 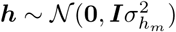. When *ρ* ≠ 0 effects of phylogenetically related haplotypes covary due to shared mutations.

For completness, the joint density of all *n* haplotype effects ***h*** is multivariate Gaussian:

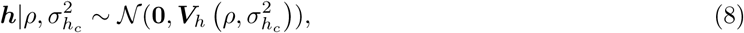

with the probability density function:

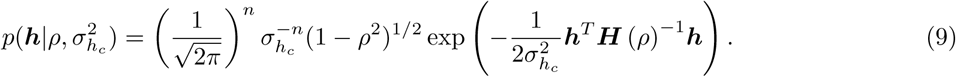

The expression (9) involves inverse of the covariance coefficient (precision) matrix ***H*** (*ρ*)^−1^, which is not necessary to compute by inverting ***H*** (*ρ*) (5). Instead, following the definition (5), inverting both sides and using the described structure of ***T*** (*ρ*)^−1^ available from the DAG and ***D*** (*ρ*), we can efficiently get this inverse by

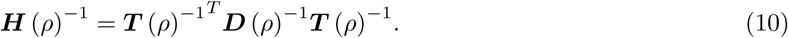

This way, we avoid computing an expensive matrix inverse.

Inspection of the structure of (10) shows that this is a very sparse matrix with a structure. We can compute the non-zero elements directly with the following simple algorithm where we loop over all haplotypes

**if** the haplotype is the “starting” haplotype **then**

add to the diagonal element 1 − *ρ*^2^

**else**

add 1 to the diagonal element

**end if**

**if** the haplotype has a parental haplotype **then**

set off-diagonal element between the haplotype and its parental haplotype to −*ρ*

add *ρ*^2^ to the diagonal element of the parental haplotype

**end if**

To fully specify the model for ***h*** (8), prior distributions must be assigned to the autocorrelation parameter *ρ* and the marginal variance 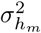 or the conditional variance 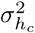. Because most mutations do not have an effect we can expect that most parent-progeny pairs of haplotypes will have similar effects, which suggests that the autocorrelation parameter will be close to 1. This knowledge can be incorporated in the prior distribution for *ρ*. For the variance parameters there may be some prior knowledge about the size of haplotype effects relative to other effects, which can also be taken into account when choosing the prior distribution. We will specify prior distributions for these parameters in later sections.

#### 2.1.3 Multiple parental haplotypes

Sometimes phylogenetic inference cannot resolve bifurcating trees with dichotomies (one parental haplotype and two progeny haplotypes) and outputs a multifurcating tree with polytomies (one parental haplotype and multiple progeny haplotypes) or even just a network (multiple parent haplotypes and multiple progeny haplotypes (e.g., Schliep et al., 2017; Uyeda et al., 2018)). We will here present an extension of the model presented in Section “The model” that can accommodate these scenarios as long as the trees or networks can be described with a DAG.

We assume that the effects of all ancestral haplotypes, the haplotypes in the top of the network, are independent and come from the same Gaussian distribution 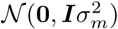. We further assume conditional independence between a haplotype and all previous haplotypes in the network given the parents of that haplotype.

In the model where each haplotype had only a single parent haplotype it was assumed that the haplotype effect was *ρ* times the parental haplotype effect plus some Gaussian noise. When a haplotype has multiple parents, we now assume that the effect is the average over each of these processes from each parental haplotype.

We illustrate this with a small example which implies the model construction used. Let haplotype *d* have parental haplotypes *a, b*, and *c*. We denote the contribution from each of these parents 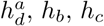, and assume

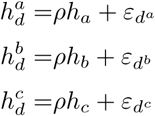

where 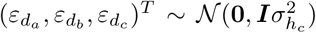. Further, we assume that the resulting effect of haplotype *h*_*d*_ is the average over all parent processes

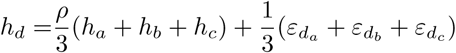

The distribution of *h*_*d*_ conditional on *h*_*a*_, *h*_*b*_ and *h*_*c*_ becomes

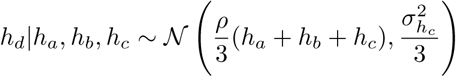

In general this means that 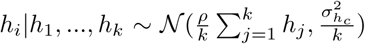, for haplotype *i* with parental haplotypes 1, …, *k*. This model construction corresponds to a model where one first takes every path down through the graph and assigns separate stationary AR(1) processes to each such path, and then assume conditionally independent but identical AR(1) processes (i.e. same parameters).

Multiple parental haplotypes change the structure of the ***T*** (*ρ*)^−1^ matrix to having −*ρ/k* value between a haplotype effect (row index) and its parental haplotype effect (column index) and ***D*** (*ρ*) matrix diagonals for “non-starting” haplotypes to 1*/k*_*i*_, where *k*_*i*_ is the number of parental haplotypes of the haplotype *i*. The algorithm to setup the ***H*** (*ρ*)^−1^ matrix is then (looping over all haplotypes)

**if** the haplotype is the “starting” haplotype **then**

add to the diagonal element 1 − *ρ*^2^

**else**

add 1*/k*_*i*_ to the diagonal element

**end if**

**if** the haplotype has a parental haplotype **then**

set off-diagonal element between the haplotype and its parental haplotype to −*ρ/k*_*i*_

set off-diagonal elements between all parental haplotypes that share a progeny haplotype to 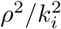

add 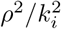 to the diagonal element of the parental haplotype

**end if**

The model presented in this section is a straightforward model, and is only one of many possible choices for a model accommodating multiple parental haplotypes.

We are only presenting one option for allowing such graph structures in the model, other choices should also be explored, which will be a topic in the discussion.

#### 2.1.4 Expanding to multiple regions due to recombination

Haplotype phylogeny can differ along genome regions due to recombination - the process of swapping genome regions between haplotypes during meiosis. We accomodate this in the haplotype network model by considering each haplotype region separately, but still within the framework of the same model. This means that the effect of haplotype *h*_*i*_ is modeled as the sum of effects for all haplotype regions. Consider haplotypes spanning three regions. The effect of haplotype *i*, is then assumed to be the sum of the effects of haplotype segments in each of the three regions:

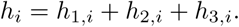

We assume the haplotype network model for each haplotype region, but with joint hyper-parameters 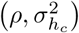. Let 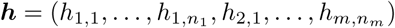 be the effect of all haplotype in all regions, where *m* is the number of regions and *n* is the number of haplotypes in each region. For ***h*** we then assume:

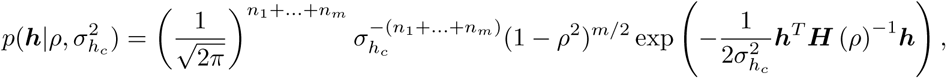

with

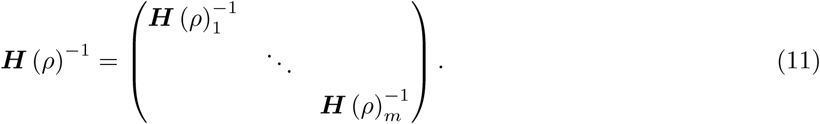

Although recombination is common, in this study we have focused on the special case of no recombination where the haplotypes are connected in one phylogeny, as presented in Section “The model”.

### 2.2 Phenotype model with haplotype effects

We show how the haplotype effects can be included in a model of phenotypic observations. We also present a phenotype model that includes independent haplotype effects or mutation effects rather than the haplotypes.

Let ***y***_*p*×1_ be phenotype observations of *p* individuals and let ***h***_*n*×1_ be the effect of *n* haplotypes obtained from phasing genotypic data of the individuals. We assume the following model for the centred and scaled phenotypic observations:

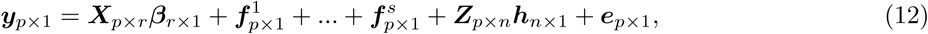

where ***β*** ∼ 𝒩 (**0, *I***1000) is a vector of *r* fixed effects with covariate matrix ***X***, 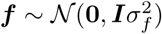 are random effects, ***h*** are the haplotype effects with incidence matrix ***Z*** that maps haplotypes to individuals, and the residual effect is 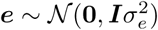. In the case of diploid individuals there will be two entries in every row of ***Z***, and a single entry for haploid individuals or male sex chromosome or mitogenome.

For the haplotype effects ***h*** we will assume three models. The first is a base model with independent haplotype effects (IH model), where 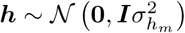. The second is the haplotype network model presented in Section “The model” (HN model), where 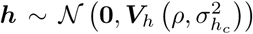. The third is an alternative way of estimating haplotype effects via a linear combination of mutation effects (mutation model). Assume ***h*** = ***Uv*** with 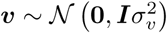 being the mutation effects and ***U*** is the matrix containing the haplotypes with reference alleles coded as 0 and alternative alleles coded as 1.

The models do not have a common intercept because a common intercept and the mean level in the haplotype effects are not identifiable when *ρ* approaches 1. Instead the mean level in the observations is captured by the haplotype effects, for computational reasons. A sum-to-zero constraint can be specified for the haplotype network part of the model if a common intercept is required, but changes the model interpretation if *ρ* is close to 1.

The identifiability problem is not special for this model, but occurs for all autoregressive models when they are used as part of a structured mixed effects model. When the goal is to make predictions about the haplotype effects, this model choice will not influence the results.

#### 2.2.1 Prior distributions

We assigned penalized complexity (PC) prior distributions to the marginal variance and the autocorrelation parameter. Penalized complexity priors are proper prior distributions developed by Simpson et al. (2017) that penalize increased complexity induced by deviation from a simpler base model to avoid over-fitting. For a random effect with a variance parameter the base models is a model where the variance of this random effect is zero. For the autoregressive model of order one we have assumed a base model with *ρ* = 1. We could have assumed a base model with *ρ* = 0, but it is more likely that phylogenetically similar haplotypes have similar effects. The penalized complexity prior can be specified through a quantile *u* and a probability *α* which satisfy Prob(*x* > *u*_*x*_) = *α*_*x*_ for the parameter *x*. Although the precision matrix is specified with the conditional variance, the prior is specified for the marginal variance. The conditions 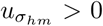 and 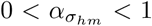 are specified for the marginal standard deviation *σ*_*hm*_. For the autocorrelation parameter we use the PC prior developed for stationary autoregressive processes (Sørbye and Rue, 2017) with base model at *ρ* = 1, and parameters satisfying −1 < *u*_*ρ*_ < 1 and 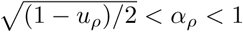.

We highlight that the prior by Sørbye and Rue (2017) was developed for a stationary autoregressive process with different model assumptions than the models presented in this paper. Ideally, the prior for the autoregressive parameter would be tailored to the haplotype network model.

### 2.3 Inference and evaluation

In this section we describe the used method for statistical inference - the Integrated Nested Laplace Approximations (INLA) - and the used methods for evaluating model fit in the simulation study.

#### 2.3.1 Inference

All models in this study fit in the framework of hierarchical latent Gaussian models, which makes INLA (Rue et al., 2009) a suitable choice to perform inference as implemented in the R (R Core Team, 2018) package INLA (available at www.r-inla.org). In this section we give a brief introduction to latent Gaussian models and how INLA is used to approximate the marginal posterior distributions in such models. For an in-depth description of INLA see Rue et al. (2009), Blangiardo and Cameletti (2015), and Rue et al. (2017).

The class of latent Gaussian models includes several models, for example generalized linear (mixed) models, generalized additive (mixed) models, spline smoothing methods, and the models presented in this article. Latent Gaussian models are hierarchical models where observations ***y*** are assumed to be conditionally independent given a latent Gaussian random field ***x*** and hyper-parameters ***θ***_1_, meaning *p*(***y***|***x, θ***_1_) ∼ Π_*i*∈ℐ_*p*(*y*_*i*_|*x*_*i*_, ***θ***_1_). The latent field ***x*** includes both fixed and random effects and is assumed to be Gaussian distributed given hyper-parameters ***θ***_2_, that is *p*(***x***|***θ***_2_) ∼ 𝒩 (***µ***(***θ***_2_), **Σ**(***θ***_2_)). The parameters ***θ*** = (***θ***_1_, ***θ***_2_) are known as hyper-parameters and control the Gaussian field and the likelihood for the data. These are usually variance parameters for simple models, but can also include other parameters, for example the *ρ* parameter in the HN model. The hyper-parameters must also be assigned prior distributions to completely specify the model.

The main aim of Bayesian inference is to estimate the marginal posterior distribution of the variables of interest, that is, *p*(*θ*_*j*_|***y***) for hyper-parameters and *p*(*x*_*i*_|***y***) for the latent field. INLA computes approximations to these densities quickly and with high accuracy. The INLA methodology is based on numerical integration of non-Gaussian hyper-paramets and utilizing Markov properties of the Gaussian parameters. Hence, for the computations to be both fast and accurate, the latent Gaussian models have to satisfy some assumptions. The number of non-Gaussian hyper-parameters ***θ*** should be low, typically less than 10, and not exceeding 20. Further, the latent field should not only be Gaussian, it must be a Gaussian Markov random field. The conditional independence property of a Gaussian Markov random field yields sparse precision matrices which makes computations in INLA fast due to efficient algorithms for sparse matrices. Lastly, each observation *y*_*i*_ should depend on the latent Gaussian field only through one component *x*_*i*_.

The R package INLA is run using the `inla()` function with three mandatory arguments: a data frame or stack object containing the data, a formula much like the formula for the standard `lm()` function in R, and a string indicating the likelihood family. Prior distributions for the hyper-parameters are specified through additional arguments. Several tools to manipulate models and likelihoods exist as described in tutorials at www.r-inla.org and the books by Blangiardo and Cameletti (2015), and Krainski et al. (2018). In the supplementary section, we have included a script showing how the data was simulated from the haplotype network model and how we fitted the model to the simulated data.

#### 2.3.2 Evaluation of model performance

We evaluated the model fit with the continuous rank probability score (CRPS) (Gneiting and Raftery, 2007). The CRPS is a proper score which takes into account the whole posterior distribution, meaning that is compares the whole estimated posterior distribution for haplotype effects with the true values and with this accounts for the uncertainty in estimation. The CRPS is negatively oriented, so the smaller the CRPS the closer the posterior is to the true value. The full Bayesian posterior output from `inla()` for these models are mixtures of Gaussians, for which there is no closed form expression for CRPS. The mixtures here are similar to plain Gaussians, so we approximate exact CRPS with the Gaussian CRPS using only the posterior mean and variances provided in the results.

We calculated CRPS for estimated haplotype effects with the IH, HN and mutation models. To ease the comparison we have then calculated a relative CRPS (RCRPS) score as the log of the ratio between the averages of the CRPS from the HN model and IH model, and correspondingly for the mutation model relative to the IH model. The score is computed as

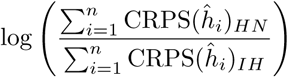

where CRPS(*ĥ*_*i*_)_*HN*_ is the CRPS of posterior haplotype effect of haplotype *h*_*i*_ with the HN model. WE will refer to this score as the RCRPS.

We also calculated the root mean square error between the mean posterior haplotype effect and true haplotype effects, but the results for the relative RMSE and RCRPS were qualitatively the same. We decided to present the RCRPS results because the CRPS takes into account the whole posterior distribution, whereas the RMSE only takes into account how closely the estimated posterior mean is to the true effect.

In addition to comparing the haplotype estimates, we compared the estimated mutation effects from the HN model and the mutation model, using the RCRPS (HN model versus mutation model). Although the HN model estimates the haplotype effects ***h***, we can obtain mutation effects via ***v*** = (***U*** ^*T*^ ***U***)^−1^***U*** ^*T*^ ***h***.

### 2.4 Simulation study

To test the proposed HN model, we first used simulated data. Here, we present data simulated from two different models − the HN model with varying degree of autocorrelation, and a more realistic mutation model where only some mutations have causal effect. We also present the models that were fitted to the simulated data, and how the model fit was evaluated.

#### 2.4.1 Simulation from the haplotype network model

We used the coalescent simulator msprime (Kelleher et al., 2016) to simulate the phylogeny shown in Figure 2 with *n* = 107 unique haplotypes. We then simulated phenotypes ***y*** for *p* = 400 individuals from the model:

**Figure 2:**
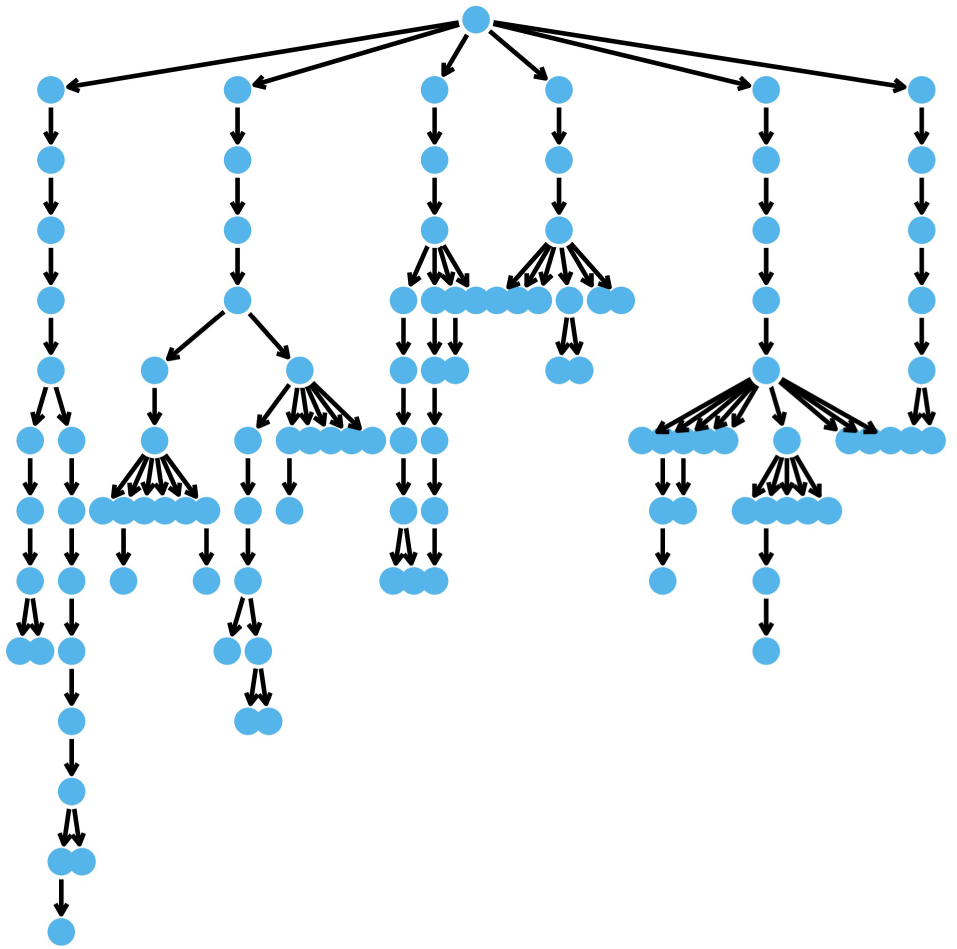
The DAG describing the phylogeny of simulated haplotypes

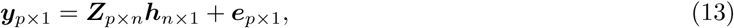

where 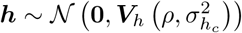 with 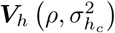 built from the DAG describing the phylogeny (Figure 2, (5)), and 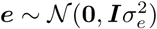.

We tested 15 parameter sets, from weak to strong haplotype effect dependency, and from low to high residual variance relative to the conditional haplotype variance:

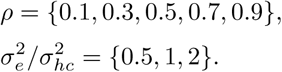

We simulated a haploid system for simplicty, so the incidence matrix ***Z*** was a zero matrix with a single 1 on each row indicating which individuals had which haplotype. We were particularly interested in estimating the haplotype effect with few or no direct phenotype observations. This is the extreme scenario where the haplotype network model could be beneficial. To achieve this, we designed the incidence matrix to create two different scenarios. In the first scenario, all haplotypes had associated phenotype observation, but some haplotypes only had one observation. We assigned a random sample of 15% of the haplotypes only to one individual each and the rest of the haplotypes randomly to the remaining individuals. In the second scenario, some haplotypes did not have phenotype observations. We selected a random sample of 15% of the haplotypes that did not have phenotype observations and assigned phenotype observations to the rest of the haplotypes.

#### 2.4.2 Simulation from the mutation model

We also simulated haplotype effects from a mutation model using the same phylogeny as in the previous section, shown in Figure 2, and using *p* = 400 individuals. For the 107 unique haplotypes we had 106 mutations in the haplotypes. We used the variants at these mutations to simulate haplotype effects and phenotypes according to the model:

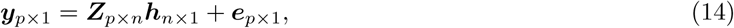

where ***h*** = ***U***_*n*×106_***v***_106×1_, ***v*** was the mutation effect, ***U*** a matrix containing ancestral (reference) alleles coded as zero and alternative alleles coded as 1, and 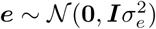. We sampled the mutation effect *v* from:

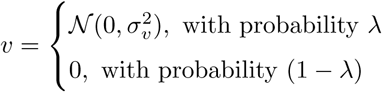

where we chose 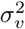 so that the empirical variance of ***h***, Var(***h***), was 1.

Again, we tested 15 parameter sets, from few to many causal variants, and from low to high residual variance relative to empirical haplotype variance

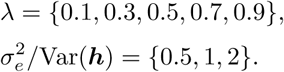

We again simulated haploid individuals, so the incidence matrix ***Z*** was a zero matrix with a single 1 on each row indicating which individuals had which haplotype. The incidence matrix was designed to create the same scenarios as for the data simulated from the HN model in Section “Simulation from the haplotype network model”.

#### 2.4.3 Models fitted to the simulated data

We fitted the HN model, IH model and the mutation model to the simulated data:

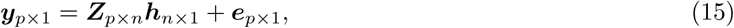

where ***h*** was assumed to be distributed according to:

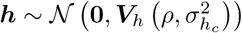 for the HN model,

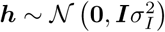for the IH model and

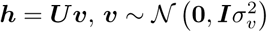 for the mutation model.

The residual effect was 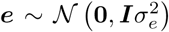. We used PC priors for the *ρ* parameters with *u*_*ρ*_ = 0.7 and *α*_*ρ*_ = 0.8, and to all variance parameters with *u* = 0.1 and *α* = 0.8.

#### 2.4.4 Evaluation

For each combination of phenotype observation distribution across haplotypes, proportion of residual variance relative to the haplotype variance and *ρ* or *λ* parameters, we performed the same experiment 50 times. In 4% of the experiments when the data was simulated from the HN model, the inference method was not able to fit the HN model and we report results only for cases where all models were successfully fitted. There was no trend for any parameter set causing the inference method to break down.

Since we created different scenarios for how phenotype observations were distributed among the haplotypes, we stratified the results for haplotype effects based on how many times a haplotype was phenotyped. For the first scenario, where some haplotypes were phenotyped either once or multiple times, we have computed the RCRPS for these two groups separately. For the second scenario, where some haplotypes were not phenotyped, we present the RCRPS only for haplotypes that were not phenotyped. In both cases, RCRPS less than zero indicates that the HN/mutation model was better than the IH model on average.

We present the RCRPS for estimated mutation effects only for the mutation model simulation, because the true mutation effects were not generated when simulating from the haplotype network model.

### 2.5 Case study: Mitochondrial haplotypes in cattle

In this section we present a case study using the haplotype network model to estimate the effect of mitochondrial haplotypes on milk yield in cattle. We first briefly describe the data and then the fitted model.

#### 2.5.1 Data

We demonstrate the use of the haplotype network model with a case study estimating the effect of mitochondrial haplotypes on milk yield in cattle from Brajković (2019). We chose this case study because mitochondrial haplotypes are passed between generations without recombination and are as such a good case for the haplotype network model. The phenotyped data comprised of information about the first lactation milk yield, age at calving, county, herd and year-season of calving for 381 cows. Additionally, the data comprised of pedigree information with 6,336 individuals (including the 381 cows) and information about mitochondrial haplotypes (whole mitogenome) variation between maternal lines in the pedigree. We inferred the mitochondrial haplotypes by first sequencing mitogenome, aligning it to the reference sqeuence and calling haplotype mutations as described in detail in Brajković (2019). We used PopART (Leigh and Bryant, 2015) to build a phylogentic network of mitochondrial haplotypes. For simplicity we used the median-joining method to show that the haplotype network model can be fit to the output of a standard phylogentic method. In this process we assumed that the ancestral alleles were the most frequent alleles. The phylogeny contained 63 unique mitochondrial haplotypes each separated by one mutation. Of the 63 haplotypes only 16 haplotypes were observed in the 381 phenotyped cows. There were five haplotypes that did not have a parent haplotype, meaning we treated them as a “starting” haplotype in the haplotype network model.

#### 2.5.2 Model

Let ***h***_*n*×1_ be the effect of the *n* = 63 mitochondrial haplotypes, and let ***y***_*p*×1_ be the phenotypes of the *p* = 381 cows. We fitted the following model to centred and scaled phenotypes:

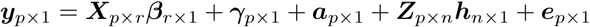

where ***β*** ∼ 𝒩 (**0, *I***1000) contained effects of age at calving as covariate and county as factor with corresponding design matrix ***X***, 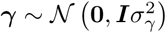 was the effect of herd-year-season of calving, 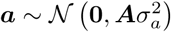 was additive genetic effect for the whole nuclear genome with the covariance coefficient matrix ***A*** derived from the pedigree (Henderson, 1976; Quaas, 1988), and lastly the mitochondrial haplotype effects were fitted with the haplotype network model 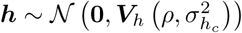 with the covariance matrix 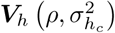 derived from the phylogeny and using the expanded model that accomodates multiple parental haplotypes from Section “Multiple parental haplotypes”. We assumed that residuals were distributed as 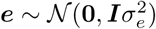.

We assigned PC priors to the *ρ* parameter with *u*_*ρ*_ = 0.7 and *α*_*ρ*_ = 0.8 and to the 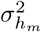 parameter with 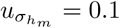 and 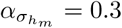, and to all remaining variance parameters with 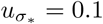 and 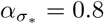.

## 3 Results

In this section we present results from the simulation study testing the behavior of the haplotype network model and the case study estimating the effect of mitochondrial haplotypes on milk yield in cattle. In the results from the simulation study, we present the RCRPS between the haploptype network (HN) model and the independent haplotype (IH) model, and between the mutation model and the IH model for the different parameter sets. In the results from the case study, we present the mean and standard deviation of the posterior mitochondrial haplotype effects mapped onto the phylogenetic network, and posterior estimates for the hyper-parameters.

### 3.1 Simulation study

#### 3.1.1 Simulation from the haplotype network model

We start by considering the results with the data simulated from the HN model from Section “Simulation from the haplotype network model” that were fitted with the models from Section “Models fitted to the simulated data”.

The RCRPS (smaller values indicate that the HN or mutation models, respectively, are better than the reference IH model) is presented in Figure 3. This figure has three panels denoting haplotypes that were observed in (A) several phenotyped individuals, (B) only one phenotyped individual and (C) were not observed in a phenotyped individual. The full lines show the RCRPS between the HN model and the IH model, while the dashed lines show the RCRPS between the mutation model and the IH model. Along the *x*-axis the autocorrelation parameter *ρ* for the simulated haplotype effects increases from weak to strong phylogenetic dependency, and the three colored lines indicate the amount of phenotypic variation due to residual relative to the variation from haplotype effects.

**Figure 3:**
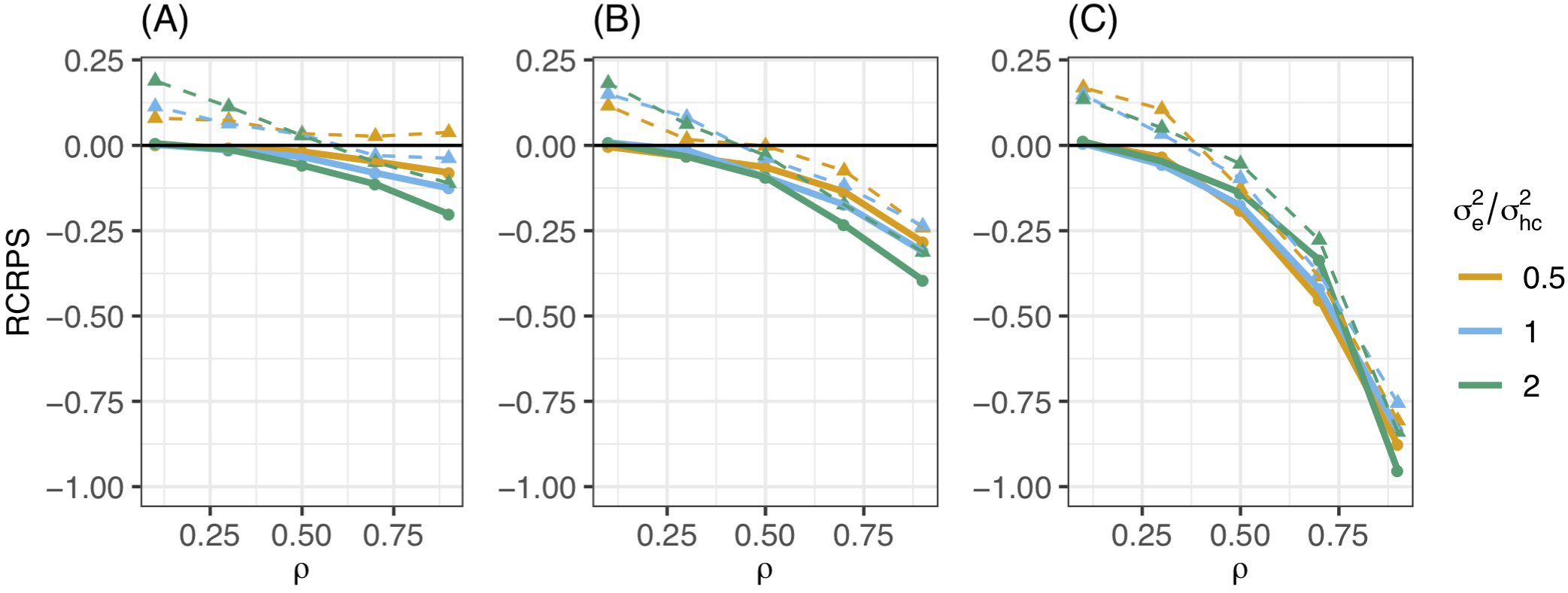
RCRPS (smaller values indicate that the HN or mutation models, respectively, are better than the reference IH model) between the HN model and the IH model (solid line) and between the mutation model and the IH model (dashed line) for data simulated from the HN model with varying *ρ* parameter and ratio between the residual and conditional haplotype variance. The three panels show RCRPS for the haplotypes that were observed in (A) several phenotyped individuals, (B) only one phenotyped individual and (C) were not observed in a phenotyped individual

In summary, Figure 3 shows that (1) the HN model outperforms the IH model across a range of model parameter values, (2) the HN model is more important for haplotypes with fewer phenotypic observations, (3) the HN model is more important for noisy phenotypic data than for phenotypic data with less noise, and (4) when haplotypes are more phylogenetically dependent, the HN model and the mutation model have similar performance. We go through each of these findings in detail.

The HN model outperforms the IH model for almost all 15 parameter sets. In all panels of Figure 3 almost all points with the full line are below zero, meaning that the HN model delivered better estimates of haplotype effects than the IH model. When the haplotype dependency due to phylogeny was low, the RCRPS was around zero, meaning that the two models were equal in estimating the haplotype effects, which was expected. As the phylogenetic dependency became stronger, the HN model improved relative to the IH model, as seen from the decreasing RCRPS as *ρ* approaches 0.9.

The improvement in CRPS with the HN model relative to the IH model increased when haplotype was observed in a smaller number of phenotyped individuals. This is indicated by the decreasing RCRPS when we compare panels (A), (B) and (C) in Figure 3, which respectively corresponds to haplotypes observed in several, one and no phenotyped individuals. The decrease in RCRPS was the largest in panel (C) followed by panel (B) and panel (A). This means that modeling phylogenetic dependency between haplotypes is most useful when there are some haplotypes with few phenotypic observations, or if we want to predict the effect of new haplotypes. Especially for haplotypes that do not have a direct link to observed phenotypes, the IH model is not useful, because it assigns the average effect of haplotypes with direct link to observed phenotypes to haplotypes without such links, whereas the HN model can assigns the haplotype effect based on a phylogenetic network. When the haplotpype effects have low phylogenetic dependency (*ρ* is low), there is not much difference in RCRPS between the three panels, as there was no similarities between the haplotype effects from which the HN model could learn from.

The improvement with the HN model relative to the IH model increased when the phenotypic data was noisier. In panels (A) and (B) in Figure 3, the RCRPS was lower with larger residual variance. This indicates that the HN model does a better separation of the environmental and genetic sources of variation than the IH model. Interestingly, we did not observe the same in panel (C), that is for haplotypes that did not have direct link to observed phenotypes. This was because the IH model performed equally poorly in predicting new haplotypes regardless of the amount of residual variance. The HN model on the other hand, performed slightly better as there was less variation due to residual effects for some values of *ρ* and similar for other values of *ρ* compared to the IH model.

As haplotypes became phylogentically more dependent with the increasing *ρ*, the HN model and the mutation model performed similarly. In all panels the dashed lines indicate a worse fit for the mutation model than for the IH model and HN model when *ρ* was low. When *ρ* increased, the mutation model improved relative to the IH model, and had a CRPS close to the CRPS for the HN model, but not better than the HN model.

#### 3.1.2 Simulated data from the mutation model

In the previous section we saw that the HN model outperformed the IH model when the simulated haplotype effects were generated from the HN model itself. Now, we consider the results with the haplotype effects simulated from a more realistic mutation model in Section “Simulation from a mutation model”, and fitted with the models from Section “Models fitted to the simulated data”. Here we varied the probability of mutations having a causal effect *λ* and we present results using only *λ* = 0.1 since the results were qualitatively similar for all tested *λ* values.

The RCRPS is presented in Figure 4 for the three different levels of phenotype observations per haplotype and three different values of residual variance relative to the empirical haplotype variance which was always 1. The full lines show the RCRPS between the HN model and the IH model, while the dashed lines show the RCRPS between the mutation model and the IH model.

**Figure 4:**
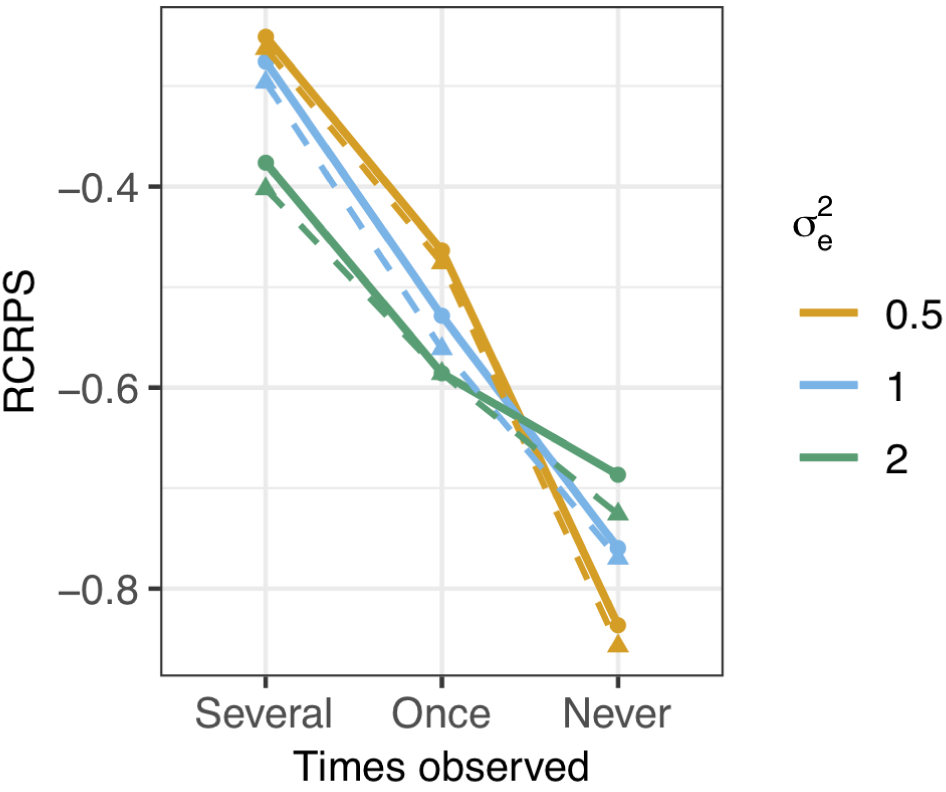
RCRPS (smaller values indicate that the HN or mutation models, respectively, are better than the reference IH model) between the HN model and the IH model (solid line) and between the mutation model and the IH model (dashed line) for data simulated from the mutation model with varying residual variance and empirical haplotype variance 1. The three scenarios show RCRPS for the haplotypes that were observed in (Several) several phenotyped individuals, (Once) only one phenotyped individual and (Never) were not observed in a phenotyped individual

In general, the results align with the results from the previous section except for the mutation model; (1) the HN model outperforms the IH model, (2) the HN model is more important for haplotypes with few phenotypic observations, (3) the HN model is more important for noisy phenotypic data and (4) the mutation model was marginally better than the HN model in estimating haplotype effects. We go through each of the findings in detail.

The HN model outperformed the IH model for all tested parameter sets. In Figure 4, all RCRPS values, are well below zero. For haplotypes observed in several or one phenotyped individual, the RCRPS was lower than what was seen in panels (A) and (B) in Figure 3. For haplotypes with no direct links to phenotype observations, the RCRPS was not improving as much as seen in panel (C) in in Figure 3.

The improvement with the HN model relative to the IH model increased with fewer phenotype observations per haplotype. The RCRPS in Figure 4 is lowest for haplotypes with no direct links to phenotype observations, second lowest for haplotypes with one direct link to a phenotype observation, and highest for haplotypes that were observed in several phenotyped individuals.

The improvement with the HN model relative to the IH model increased with increasing residual variation. In Figure 4 the RCRPS for haplotypes observed in several or one phenotyped individual decreases with increasing residual variance. This was again not the case for haplotypes with no direct links to phenotype observations. As mentioned in the previous section, the IH model is predicting new haplotypes equally poorly irrespective of the residual variance. The HN model on the other hand, improves the prediction of new haplotypes when the phenotypic data is less noisy. This explains why the improvement in CRPS between the HN model and IH mode for haplotypes with no direct links to phenotype observations is largest when with less residual variation, and why the RCRPS in panel (C) is lowest for residual variance 0.5.

The mutation model was marginally better than the HN model in estimating haplotype effects. The dashed lines in Figure 4 indicate the RCRPS between the mutation model and the IH model, and the full lines indicate the RCRPS between the HN model and the IH model. The dashed lines and full lines follow each other closely, and the dashed lines are slightly lower than the full lines, indicating that the mutation model was slightly better than the HN model, although not by much.

In Table 2 we present the average RCRPS between the HN model and the mutation model for the estimated mutation effects. This table has the average RCRPS and its standard error for the two scenarios where either all haplotypes had associated phenotype observation, or most haplotypes had associated phenotype observation and the rest did not, with different proportions of mutations with causal effect and for different residual variance. Averages above zero indicates that the mutation model had better CRPS, and averages below zero indicates that the HN model had better CRPS. Overall the difference between the two models is small. The mutation model had the best performance when there were few causal mutations, and the HN had the best performance when there were many causal mutations. This corresponds with the results presented for the estimated haplotype effects in Figure 4. The difference between the two models decreased as the proportion of mutations with causal effect out of all mutations increased.

**Table 2:**
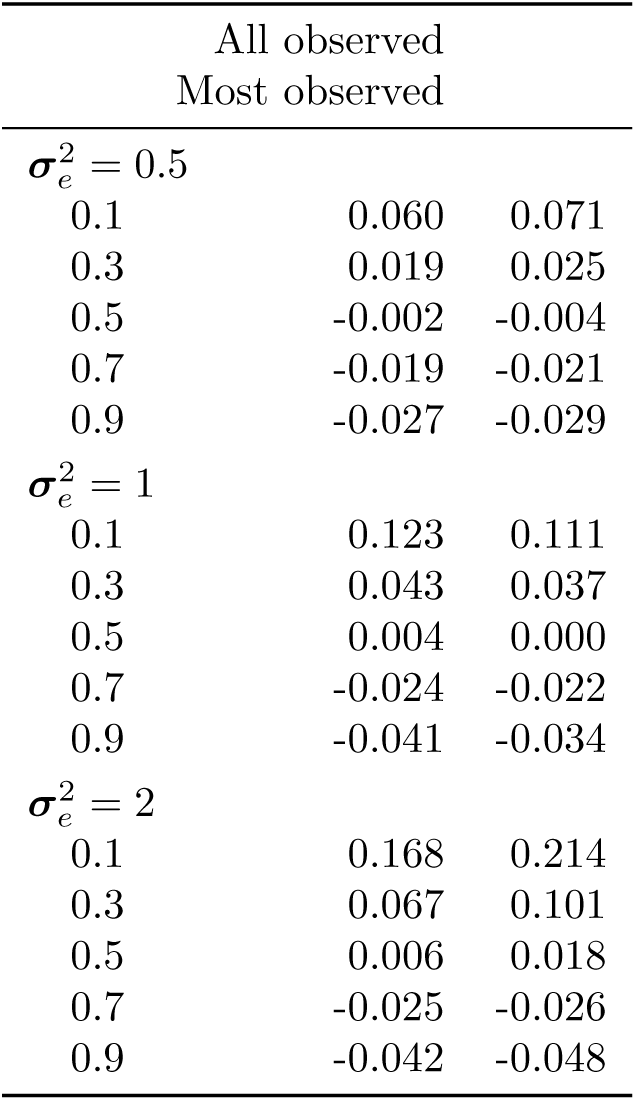
RCRPS between the HN model and the mutation model for mutation effects by different values of residual variance, proportion of causal mutations and for the two scenarios where either all or most haplotypes have direct links to observed phenotypes

|  | All observed |  |
| --- | --- | --- |
|  | Most observed |  |
| $\sigma_e^2 = 0.5$ | | |
| 0.1 | 0.060 | 0.071 |
| 0.3 | 0.019 | 0.025 |
| 0.5 | -0.002 | -0.004 |
| 0.7 | -0.019 | -0.021 |
| 0.9 | -0.027 | -0.029 |
| $\sigma_e^2 = 1$ | | |
| 0.1 | 0.123 | 0.111 |
| 0.3 | 0.043 | 0.037 |
| 0.5 | 0.004 | 0.000 |
| 0.7 | -0.024 | -0.022 |
| 0.9 | -0.041 | -0.034 |
| $\sigma_e^2 = 2$ | | |
| 0.1 | 0.168 | 0.214 |
| 0.3 | 0.067 | 0.101 |
| 0.5 | 0.006 | 0.018 |
| 0.7 | -0.025 | -0.026 |
| 0.9 | -0.042 | -0.048 |

### 3.2 Case study: Mitochondrial haplotypes in cattle

In this section we present results for the case study of estimating the effect of mitochondrial haplotypes on milk yield in cattle presented in Section “Case study: Mitochondrial haplotypes in cattle”. We present the posterior mean and standard deviation for the effect of mitochondrial haplotypes mapped onto the phylogeny, the posterior distribution for the autocorrelation parameter *ρ*, and the mean and 95% confidence interval of the posterior variances in the model.

In summary the results show that there was (1) sharing of information between the mitochondrial haplotypes, (2) haplotypes without direct link to observed phenotyopes were estimated with larger uncertainty, (3) indications of strong phylogenetic dependency between the haplotypes and (4) significant, proportion of the total phenotypic variation explained by mitochondrial haplotypes. We now go through each of these findings in detail.

The HN model enabled sharing of information from the haplotypes that had a direct link with observed phenotypes to the other haplotypes. In Figure 5 we present the posterior mean for the effect of mitochondrial haplotypes with node color. Haplotype effect estimates are similar for phylogenetically similar haplotypes, meaning that there was sharing of information between the haplotypes. The figure also shows that the few haplotypes that had direct links with phenotype observations (nodes labeled with 1) were separated from the other haplotypes with a substantial number of mutations yet the information was shared through the phylogeny to all haplotypes, which was the aim of the HN model.

**Figure 5:**
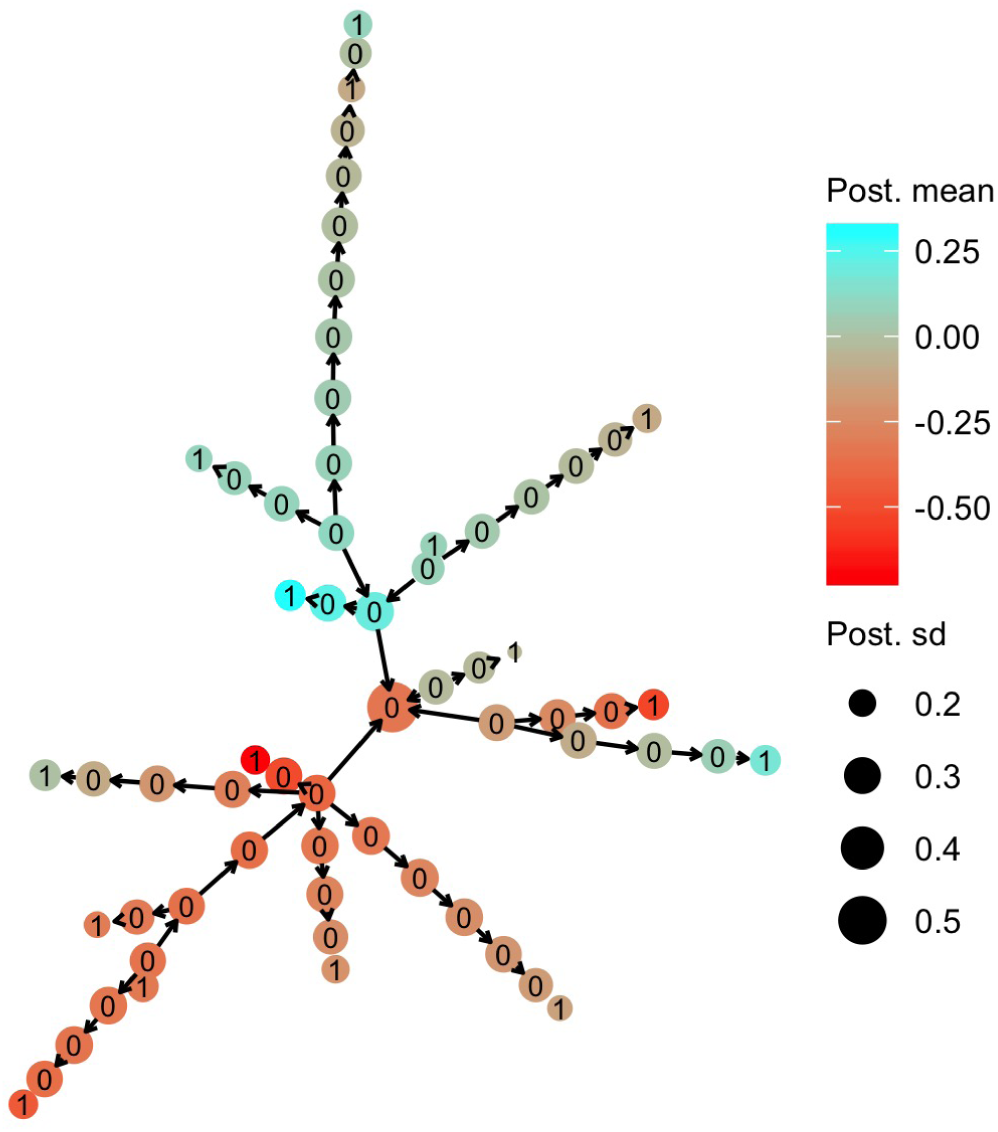
Posterior mean and standard deviation for mitochondrial haplotype effects on milk yield in cattle. Posterior means are denoted with node color, while posterior deviations are denoted by the node size. The numbers on each haplotype node indicate if the haplotype had a direct link to the observed phenotyope (1) or not (0)

Haplotypes without direct links to observed phenotypes were estimated with larger uncertainty. In Figure 5 we present the posterior standard deviation for the effect of mitochondrial haplotypes with node size. We see that the haplotypes with direct links to observed phenotypes (nodes labeled with 1) have smaller posterior standard deviation than the other hapolotypes (nodes labeled with 0). The posterior standard deviation decreased slightly as the haplotypes without direct links were closer (in number of mutations) to the haplotypes with direct links, which was expected. However, the overall posterior standard deviations for haplotype effects were relatively large, because the data set was small and there were few haplotypes with direct links to observed phenotypes connected to the other haplotypes with many mutations between them.

The posterior distribution for the autoregression parameter *ρ* indicated strong dependency between haplotype effects. The posterior distribution of *ρ* is shown in Figure 6. The mode of the distribution lies around 0.85, and the mean lies around 0.73, indicating that neighboring haplotypes had similar effects, which is related to the sharing of information between haplotypes seen in Figure 5.

**Figure 6:**
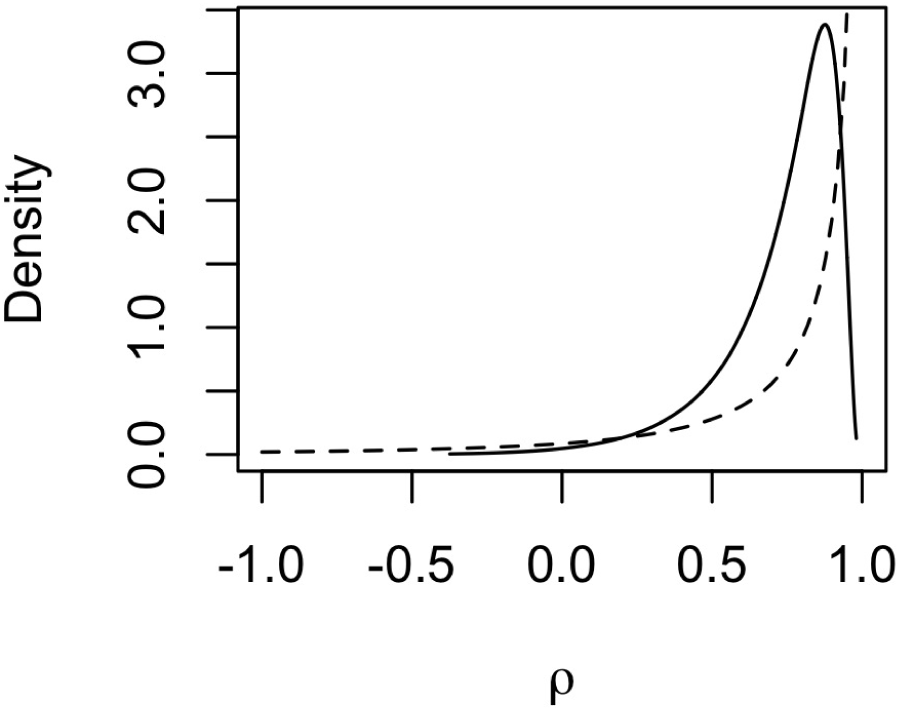
Prior (dashed line) and posterior (solid line) distribution for the autocorrelation parameter *ρ* for mitochondrial haplotype effects on milk yield in cattle

A significant amount of the total phenotypic variation was explained by the mitochondrial haplotypes. In Table 3 we present the posterior mean and 95% confidence interval of each variance component in the model, and how much of the total variation in the model 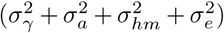 was explained by each variance component. The posterior distribution of the conditional haplotype variance was obtained by computing 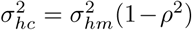, using 10000 samples from the posterior distributions of the marginal haplotype variance and the autocorrelation parameter. We see that the marginal haplotype variance 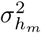 and conditional haplotype variance 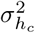 is smaller compared to the additive genetic variance 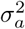, and the residual variance 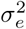. This was expected as the mitogenome (∼ 1 × 16*Kbp*) is much smaller than the nuclear genome (∼ 2 × 3*Gbp*). In the light of this difference we can say that mitochondrial haplotypes captured a significant amount of phenotypic variation. The variance for the random effect of herd-year-season of calving 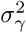 was also smaller compared to 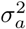 and 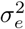.

**Table 3:**
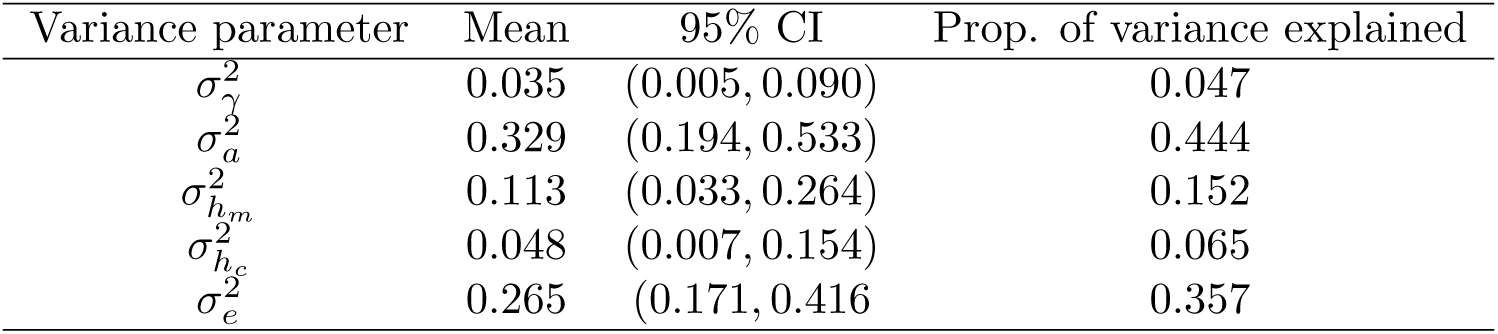
Posterior mean, 95% confidence interval (CI) for variance parameters, and the proportion of variation explained by each variance component for the case study estimating mitochondrial haplotype effects on milk yield in cattle

| Variance parameter | Mean | 95% CI | Prop. of variance explained |
| --- | --- | --- | --- |
| $\sigma_{\gamma}^2$ | 0.035 | (0.005, 0.090) | 0.047 |
| $\sigma_a^2$ | 0.329 | (0.194, 0.533) | 0.444 |
| $\sigma_{h_m}^2$ | 0.113 | (0.033, 0.264) | 0.152 |
| $\sigma_{h_c}^2$ | 0.048 | (0.007, 0.154) | 0.065 |
| $\sigma_e^2$ | 0.265 | (0.171, 0.416) | 0.357 |

It should be noted that this is a small dataset with few haplotypes with direct links to observed phenotypes, which causes the posterior estimates to be strongly influenced by the prior distributions, especially the posterior for *ρ*. However, we still chose to assign an informative prior to *ρ*, since it is expected that most mutations have no causal effect.

## 4 Discussion

The objective of this paper was to propose a hierarchical model that leverages haplotype phylogeny to improve the estimation of haplotype effects. We have presented the haplotype network model, evaluated it using simulated data from two different generative models, and applied it in a case study of estimating the effect of mitochondrial haplotypes on milk yield in cattle. Here, we highlight three points for discussion in relation to the proposed haplotype network model: (1) the importance of the haplotype network model, (2) future development and possible extensions and (3) limitations.

### 4.1 The importance of the haplotype network model

We see three important advantages of the haplotype network model. These are specifcally the ability to share information between related haplotypes, computational advantages when modeling a single region of a genome and the potential to capture background specific mutation effects.

The haplotype network model utilizes phylogenetic relationships between haplotypes and with this improves estimation of their effects. From the simulation study, we saw the importance of this information sharing when there is limited information per haplotype. For example, we were able to estimate the effect of haplotypes that had few or no direct links to observed phenotypes with much higher accuracy than with a model assuming independent haplotypes. In the haplotype network model the autocorrelation parameter *ρ* and the conditional variance parameter 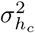 reflect the effects of phylogenetically similar haplotypes. As the autocorrelation approaches 1, haplotype effects become more dependent. Further, if conditional variance is small the large dependency and small deviations lead to similar effects for phylogenetically similar haplotypes, suggesting that mutations separating the haplotypes have very small or no effect compared to other shared mutations between haplotypes. If on the other hand conditional variance is large, the large dependency and large deviations lead to haplotype effects that change rapidly along the phylogeny, suggesting that mutations separating the haplotypes have large effects. On the other hand, if the autocorrelation parameter approaches 0, the dependency between phylogenetically similar haplotypes is decreasing, suggesting that modelled mutations do not have an effect and that haplotypes should be modelled independently.

The three extreme scenarios of hyper-parameter values could denote three real cases. The first case with high autocorrelation and small conditional variance could reflect a situation where the whole haplotype sequence would be used to build phylogeny and since most mutations do not have a causal effect, but some do, it is expected that similar haplotypes will have similar effects with small differences between the haplotypes. The second case with high autocorrelation and large conditional variance could reflect the situation when the number of causal mutations would be high compared to all mutations (because only such mutations are analysed) and therefore change of effects along the phylogeny would be larger. The third scenario with no autocorrelation could reflect the situation where phylogeny does not correlate with phenotype change, which can be due to many reasons (analysis of a genome region that is not associated with the phenotype, inadequate genomic platform to capture genotype - phenotype association, etc.).

As mentioned in the introduction, modeling phenotypic variation as a function of haplotype variation has extensive literature (Templeton et al., 1987; Balding, 2006; Thompson, 2013; Morris and Cardon, 2019). The prime motivation for this work is the recent growth in the generation of large scale genomic datasets and methods to build phylogenies (Kelleher et al., 2019). To this end we aimed to develop a general haplotype network model that could exploit phylogenetic relationships between haplotypes in a computationally efficient way. Namely, the model uses phylogeny encoded with a DAG, which enables sharing of information between similar haplotypes in a recursive way that also implies computational benefits. The computational benefits come from the sparse precision matrix 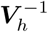, which is due to the conditional independence structure encoded in the DAG of a network of haplotypes (Rue and Held, 2005). Sparsity is important, because it enables fitting large models due to smaller memory requirement and faster calculations (Rue and Held, 2005). The computational benefits are not critical when the number of haplotypes is small. In that case the matrix ***V***_*h*_ is small and easy to invert, though for the autoregressive model we would have to invert it many times during the estimation procedure due to dependency on the autocorrelation parameter. However, it is better to avoid inversions if possible because it can lead to numerical errors and loss of precision (e.g., Misztal, 2016).

While the haplotype network model is different to the pedigree mixed model (Henderson, 1976; Quaas, 1988) (where we model the inheritance of whole genomes in a pedigree without (fully) observing the genomes) or the phylogenetic mixed model (Lynch, 1991; Pagel, 1999; Housworth et al., 2004; Hadfield and Nakagawa, 2010) (where we model the inheritance of whole genomes in a phylogeny without (fully) observing the genomes), the principles of conditional dependence between genetic effects and the resulting sparsity are the same (Rue and Held, 2005). The key difference of the haplotype network model is that it estimates the effect of observed haplotype sequences as compared to unobserved or partially observed inheritance of genomes in a pedigree or phylogeny. To improve the estimation of the haplotype effects we take into account the phylogenetic relationships. While the use of phylogenetic relationships might seem redundant if we know (most of) the haplotype sequence, the simulations showed that it improves estimation in most cases, even marginally compared to the mutation model where we directly model mutation effects. The haplotype network model can be seen as a hybrid between the mutation model (that models variation between the columns of a haplotype matrix) and the independent haplotype model (that models variation between the rows of a haplotype matrix). This hybrid view might improve genome-wide association studies (see review by Gibson, 2018; Simons et al., 2018; Uricchio, 2019; Morris and Cardon, 2019).

Further, the haplotype network model has the potential to capture background specific mutation effects. Background specific mutation effects are observed when the effect of a mutation depends on other mutations present in an individual (e.g., Chandler et al., 2017; Wojcik et al., 2019; Steyn et al., 2019). Such effects can also manifest when a mutation is marking another unobserved mutation with correlation that varies between backgrounds. The haplotype network model can capture background specific mutation effects through the fact that it is modeling haplotype effects and not mutation effects. If there are background specific mutation effects the haplotype effect differences will capture this, while a mutation model only estimates an average effect of a mutation across multiple backgrounds (haplotypes). We must point though that the haplotype network model captures only local effects, that is due to interactions between mutations present on a haplotype (e.g., Clark, 2004; Liu et al., 2019). We have not evaluated how well the model captures background specific mutation effects in this study and more simulations and applications to a range of datasets are needed to evalute this aspect.

### 4.2 Future development and possible extensions

There is a number of areas for future development with the haplotype network model. We are looking into four areas: making the model more flexible in the number of mutations separating phylogentically similar haplotypes, modeling haplotype differences in a continuous way utilising branch lengths, incorporating biological information and phylogenetic aspects of haplotype relationships.

We have developed the haplotype network model by assuming the differences between similar haplotypes is due to one mutation to simplify model definition. However, in the observed data there might not be haplotypes that are separated for just one mutation. We handle this situation by inserting phantom haplotypes. The order of mutations in such situations is uncertain and a model could be generalized to account for these larger number of mutations between haplotypes. However, the current “one-mutation” difference model setup has a useful property of inferring the value of unobserved haplotypes and the sparse model definition does not increase computational complexity of the model.

Relatedly, the haplotype network model could be generalized to utilize time calibrated distances between haplotypes rather than using the number of mutations. The OrnsteinUhlenbeck (OU) process is the continuous-time analogue of the autoregressive process of order one used in this study, and plays a major role in the analysis of the evolution of phenotypic traits along phylogenies (Lande, 1976; Hansen and Martins, 1996; Martins and Hansen, 1997; Paradis, 2014). Relatedly, if the autocorrelation parameter of the autoregressive process of order one is set to 1 we get the non-stationary discrete random walk process, whose continuous-time analogue is the Brownian process that is the basic model of phylogenetic comparative analysis (Felsenstein, 1988; Huey et al., 2019). The haplotype network model here uses a parameterization that only allows non-singular models, which rules out pure random walk processes, but a singular version with *ρ* = 1 can also be formulated and implemented in R-INLA. The merit of one model over the other will depend on specific datasets and genome regions as well as the scope of analysis (macroevolution vs. microevolution of a genome region). There is a scope to improve computational aspects for these continuous models too by employing recent developments from the statistical analysis of irregular time-series (Lindgren and Rue, 2008).

In the haploptype network model presented in this study, the same autocorrelation parameter has been assumed for all mutations. However, the autocorrelation parameter could be allowed to vary as Beaulieu et al. (2012) did in the context of adaptive evolution. The stationary autoregressive process of order one for trees with only one ancestral haplotype and no recombination allows for such extensions without having to change the variance parameter. For example, one could use different autocorrelation parameters for different types of mutations to incorporate biological information into the model. This would enable combining the quantitative analysis of mutation and haplotype effects from this study with molecluar genetic tools such as Variant Effect Predictor (McLaren et al., 2016).

In this study we have assumed that the phylogenetic network is given and described with a DAG. There is a large body of literature on inferring phylogenies in the form of strict bifurcating trees, more general trees or networks and recent developments in genomics are rapidly advancing the field (e.g., Anisimova, 2012; Puigbò et al., 2013; Schliep et al., 2017; Uyeda et al., 2018). We have named the model the haplotype network model, because it can work both with phylogenetic bifurcating and multifurcating trees and phylogenetic networks. The only condition is that we describe the haplotype relationships with a DAG, which gives the structure to the hierarchical haplotype model. Many tools provide such output (e.g., Leigh and Bryant, 2015; Suchard et al., 2018; Kelleher et al., 2019). To accomodate general DAGs where a haplotype node could potentially have multiple parental haplotype nodes we have generalized the model construction to allow for network structures. This generalisation also enables the model to describe haplotype relationships without paying attention to the directionality as long as there are no directed loops in the graph.

Knowing the order of mutations and therefore which haplotypes are parental to other haplotypes is beneficial because it leads to a tree structure and a sparser model (Rue and Held, 2005). An example of non-optimal sparsity can be seen in our case study. In Figure 5 the “central” haplotype with the largest uncertainty is modelled as a progeny haplotype of four surrounding haplotypes, which means that there is a dense 5×5 block in the precision matrix 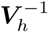. The block is dense because the “central” haplotype is modelled as a function of the other four “parental” haplotypes, which invokes conditional dependence between the “parental” haplotypes. If however the “central” haplotype would have been used as the parental haplotype the 5×5 block would be sparse since all other haplotypes would be conditionally independent given the “central/parental” haplotype. The same applies also for the other parts of the haplotype network in Figure 5. These examples of non-optimal sparsity are a consequence of the haploptype network we used, but we did this for simplicity and to emphasize the flexibility of the haploptype network model.

The haploptype network model could also work with probabilistic networks where edges have associated uncertainty (weights). If we can encode such a network with a directed acyclic graph then the edge weights can be used in model construction - for example, in the same way uncertain parentage is handled in pedigree models (Henderson, 1976). An alternative would be to construct a model for each possible realisation of a network, run separate models and combine haplotype estimates in the spirit of Bayesian model averaging. This latter approach is obviously computationally more demanding.

### 4.3 Limitations

The haplotype network model also has some limitations that merit further development. We highlight three areas: is the haplotype network model necessary given that we can model mutation effects, Gaussian assumption and causal mutations, and modeling recombining haplotypes.

For the haplotype network model to achieve its full potential, the data needs to have a certain structure. We saw from fitting the hapolotype network model to a real data set, that having few haplotypes with direct links to observed phenotypes and many haplotypes without, meant that we had large uncertainty in estimated haplotype effects. We also saw from the simulated data, that the mutation model was slightly better at estimating the mutation effects than the haplotype network model, when the data was simulated from a mutation model, but the magnitude of difference was minimal. In the future, different data structures with balanced and unbalanced structure spanning multiple populations with varying levels of connectedness, small or large number of mutations, and causal or non-causal mutations should be tested to find optimal scenarios for the haplotype network model to achieve its full potential.

The haplotype network model assumes that the haplotype effects follow a Gaussian distribution. If all, or very many, of the haplotypes have the same effect, the true haplotype effect distribution may be quite different from Gaussian, which breaks the model assumptions and perhaps other models should be proposed. Blomberg et al. (2019) describe the underlying theory behind the common Gaussian processes, such as Brownian motion and Ornstein-Uhlenbeck process, and present general methods for deriving new stochastic models, including new non-Gaussian models of quantitative trait macroevolution. See also Schraiber and Landis (2015); Landis et al. (2012); Duchen et al. (2017). Application of these models will depend on the magnitude of deviations from Gaussian assumptions, which might be large on the scale of macroevolution, but might also be large when looking at a specific genome region.

Scaling the haplotype network model to multiple recombining haplotype regions is challenging for two reasons. First, while phasing methods have improved substantially in the last years (Marchini, 2019), determining a recombination breakpoint across multiple individuals is abritrary. Second, the sparsity of the haplotype network model comes from the sparsity of the precision matrix 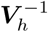, which is part of the prior distribution for haplotype effects. When developing extension for recombining haplotypes we observed that the sparsity in the prior is maintained also for multiple consequitive haplotype regions along a chromosome as shown in (11). However, we observed that the design matrices that link phenotype observations with multiple haplotype regions start to create dense cross-products in the normal system of equations as we increase the number of regions and the sparsity advantage from one haplotype region is lost. To this end we are exploring alternative ways of formulating the haplotype network model following Kelleher et al. (2019). Further research is needed to be able to scale the haplotype network model to many haplotype regions or even whole chromosomes and genomes.

## Author Contributions

MLS, FL and GG conceived and derived the haplotype network model. MLS, IS, and GG designed the analysis, and evaluated the results. MLS simulated data and performed all analyses. VB and VCC provided case study data. MLS wrote the manuscript, and FL, IS and GG commented on and edited the manuscript. All authors have read and approved the final manuscript.

## Data Availability Statement

Simulation of data and its analysis is possible through simulation code. The case study data on mitochondrial haplotypes and phenotype data in is available on research collaboration basis from Vladimir Brajkovic and Vlatka Cubric-Curik. This data was used only to demonstrate the use of the haplotype network model and have not been used to draw conclusions supporting the findings.

